# Enhanced Suppression of *Stenotrophomonas maltophilia* by a Three-Phage Cocktail: Genomic Insights and Kinetic Profiling

**DOI:** 10.1101/2024.08.14.607921

**Authors:** Alisha N. Monsibais, Olivia Tea, Pooja Ghatbale, Jennifer Phan, Karen Lam, McKenna Paulson, Natalie Tran, Diana S. Suder, Alisha N. Blanc, Cyril Samillano, Joy Suh, Sage Dunham, Shane Gonen, David Pride, Katrine Whiteson

## Abstract

In our era of rising antibiotic resistance, *Stenotrophomonas maltophilia* (STM) is an understudied, gram-negative, aerobic bacterium widespread in the environment and increasingly causing opportunistic infections. Treating STM infections remains difficult, leading to an increase in disease severity and higher hospitalization rates in people with Cystic Fibrosis (pwCF), cancer, and other immunocompromised health conditions. The lack of effective antibiotics has led to renewed interest in phage therapy; however, there is a need for well-characterized phages. In response to an oncology patient with a respiratory infection, we collected 18 phages from Southern California wastewater influent that exhibit different plaque morphology against STM host strain B28B, cultivated from a blood sample. Here, we characterize the genomes and life cycle kinetics of our STM phage collection. We hypothesize that genetically distinct phages give rise to unique lytic life cycles that can enhance bacterial killing when combined into a phage cocktail compared to the individual phages alone. We identified three genetically distinct clusters of phages, and a representative from each group was screened for potential therapeutic use and investigated for infection kinetics. The results demonstrated that the three-phage cocktail significantly suppressed bacterial growth compared to individual phages when observed for 48 hours. We also assessed the lytic impacts of our three-phage cocktail against a collection of 46 STM strains to determine if a multi-phage cocktail can expand the host range of individual phages. Our phages remained strain-specific and infect >50% of tested strains. The multi-phage cocktail maintains bacterial growth suppression and prevents the emergence of phage-resistant strains throughout our 40-hour assay. These findings suggest specialized phage cocktails may be an effective avenue of treatment for recalcitrant STM infections resistant to current antibiotics.

**IMPORTANCE:** Phage therapy could provide a vital strategy in the fight against antimicrobial resistance (AMR) bacterial infections; however, significant knowledge gaps remain. This study investigates phage cocktail development for the opportunistic pathogen *Stenotrophomonas maltophilia* (STM). Our findings contribute novel phages, their lytic characteristics, and limitations when exposed to an array of clinically relevant STM strains. Eighteen bacteriophages were isolated from wastewater influent from Escondido, California, and subjected to genomic analysis. We investigated genetically distinct phages to establish their infection kinetics and developed them into a phage cocktail. Our findings suggest that a genetically distinct STM phage cocktail provides an effective strategy for bacterial suppression of host strain B28B and five other clinically relevant STM strains. Phage therapy against STM remains poorly understood, as only 39 phages have been previously isolated. Future research into the underlying mechanism of how phage cocktails overwhelm the host bacteria will provide essential information that could aid in optimizing phage applications and impact alternative treatment options.

## INTRODUCTION

Antimicrobial resistance (AMR) in a clinical setting occurs when infecting microbes overcome antimicrobial medication, ultimately leading to severe disease and mortality in the infected patient. By 2050, AMR is projected to contribute to over 10 million deaths annually^1^, leading to an economic impact of $300 billion as treatment will be prolonged and less effective^2^. This impending crisis has been connected to the misuse and overuse of antibiotics in the clinical setting and agriculture industry^3^. Reduced investment in antibiotic discovery has also intensified AMR infection rates and impacts^4,5^. However, one approach with the potential to mitigate these hard-to-treat recalcitrant infections is phage therapy^6^.

Phage therapy utilizes lytic bacteriophages, viruses that infect bacteria, to reduce bacterial burdens associated with infections^7^. Phages attach to the host bacterial cell via specific receptors, inject phage DNA, and hijack host machinery, ultimately resulting in the host cell death by lysis and progeny virus release^8,9^. Although this knowledge of phage biology has been around for a century, basic research into phage safety, antibacterial properties, and best practices for therapeutic use have been understudied^10,11^. However, with the rise in AMR infections and the increased use of therapeutic phages, basic phage biology has taken on new importance. Indeed, phage therapy has shown promising results in life-threatening infections in various multidrug-resistant (MDR) bacteria^6,12,13^, and clinical trials are currently underway^14,15^.

Developing safe and effective phages for therapy will benefit significantly from a thorough characterization of phages, especially in their infection kinetics. Screening of phage candidates begins with genomic sequencing to assess the presence of AMR and toxin-ending genes, which would exclude the phage from use^11^. Determining infection kinetics includes tracking the rate of phage attachment to its host cell^16^ and tracking the life cycle of the phage via a one-step growth curve^17^, which measures the length of the latent phase, burst size, and duration of phage infection. Both of these time-dependent phage-bacteria interactions are important in identifying the underlying phage selection pressures and antibacterial properties, which may aid in strengthening phage therapy treatment options.

Current data suggest individual phages generally have a narrow host range, meaning they can only infect a subset of strains from a single bacterial species^7,18^. Since phage and bacteria co-evolve in response to one another, using multi-phage cocktails has enhanced the lytic outcomes of MDR bacteria^19,20^. Bacteria resist phage through several mechanisms, including restriction-modification systems, CRISPR-Cas9 immunity, and abortive infection^21–23^. Thus, when host bacteria are exposed to single phages, previous data has shown resistance can quickly arise, emphasizing the need for phage cocktails, which may mitigate the development of resistance^24,25^. Indeed, prior work in our and other laboratories has demonstrated that cocktails increase phage infectivity by reducing the growth of the target pathogen and limiting the development of phage resistance^24,26,27^. Thus, designing cocktails is an essential aspect of improving the efficiency of phage therapy.

*Stenotrophomonas maltophilia* (STM) is a gram-negative emerging opportunistic pathogen that has plagued immunocompromised individuals and people with cystic fibrosis^28^. STM is innately antibiotic-resistant, containing an extensive repository of AMR mechanisms such as biofilm formation and beta-lactamases^29,30^. Additionally, clinical isolates have higher mutation rates than their environmental counterparts, enabling them to adapt quickly^31^. A recent meta-analysis of STM global prevalence revealed an increased trend of STM infections over the last 30 years, along with increased antibiotic resistance in both tigecycline and ticarcillin-clavulanic acid^32^. Thus, there is a clear need to investigate phages against STM, considering only 39 phages have been isolated against this opportunistic pathogen, and no phage cocktail studies have been reported as of this writing^33–41^.

We hunted for phages in Southern California sewage influent and ultimately found 18 phages that could infect an STM strain isolated from an oncology patient’s blood sample. We used these phages to address the following questions: (1) How genetically diverse are these 18 phages? (2) What are the phage infection kinetics of genetically unique STM phages? (3) Can a phage cocktail comprising several genetically unique phages extend the lytic activity of the phages and suppress bacterial growth? We hypothesized that genetically distinct phages would give rise to unique lytic life cycles, which can enhance lytic activity when combined into a phage cocktail compared to the individual phages alone.

## RESULTS

### Comparative genomic analysis and bioinformatic screening of STM phages

We isolated 18 phages from Southern California wastewater influent against STM strain B28B, a bacterial isolate from an oncology patient (**Supp Table 1 and 2**). Coverage analysis was conducted on each phage to ensure adequate coverage of sequencing reads (**Supp Fig. 1**), and CheckV analysis was used to determine the completeness of the genome (**Supp Table 3**). The average nucleotide identity percentage (ANI%) of the 18 phages in our STM phage collection revealed three distinct phage clusters and a singleton isolate (**Figure 1A**), which was confirmed with VIRIDIC analysis (**Supp Fig. 2**). Additional comparative genomics visualization using ANVI’O shows the gene clusters organized in a similar pattern, based on genetically distinct cohorts (**Supp Fig. 3**). Additionally, BLASTn analysis was conducted on each phage to assess similarity to previously identified phages (data not shown)^42^. The top right cluster (**Figure 1A**) contained a high degree of similarity (>98% ANI) and a siphovirus morphology was indicated by collective BLASTn hits to Caudoviricetes sp. isolate 94, Caudoviricetes sp. isolate 231, Caudoviricetes sp. isolate 163, *Stenotrophomonas* phage CUB19 and Siphoviridae environmental samples clone NHS-Seq1. The bottom left phage cluster (**Figure 1A**) also contained variation in similarity with 86-99 ANI%, and a podovirus morphology was indicated by collective BLASTn hits to *Stenotrophomonas* phage Ponderosa, *Stenotrophomonas* phage Ptah, *Stenotrophomonas* phage Pepon, and *Stenotrophomonas* phage TS-10. Phage ANB28 was a stand-alone phage isolate, and BLASTn analysis demonstrated that it was 73.88% similar to *Xanthomonas* phage JGB6, though the phage morphology was unknown. After comparative genomic analysis, we selected one representative from each group: ANB28, KB824 (podovirus), and SBP2ɸ2 (siphovirus). Phylogenetic analysis was performed using our three distinct phages against 27 previously discovered STM phages using ViPtree, a program used to generate viral proteomic trees based on genome-wide similarities derived from tBlastx^43^. ANB28 and SBP2ɸ2 diverge from previously isolated STM phages, while KB824 is closely related to *Stenotrophomonas* phage Ponderosa, consistent with BLASTn results (**Figure 1B**). Bioinformatic screening of the genomes from the three representative phages revealed no genome-encoded integrase, AMR, or toxin genes (**Table 1**). ANB28 had the largest genome at 108 kb, which consisted of 194 open reading frames (ORFs) and five tRNAs. KB824 had the shortest genome at ∼43 kb, which consisted of 76 ORFs and zero tRNAs. SBP2ɸ2 had a genome of ∼50 kb, which consisted of 123 ORFs and eight tRNAs. Annotations of gene maps for each phage were created by listing genes with predicted annotations on the top row, unlabeled hypothetical proteins on the bottom row, and tRNAs denoted in green located on the genome line (**Figure 2**). These phylogenetic and genomic results confirm that our three selected phages are genetically distinct.

**Figure 1.**
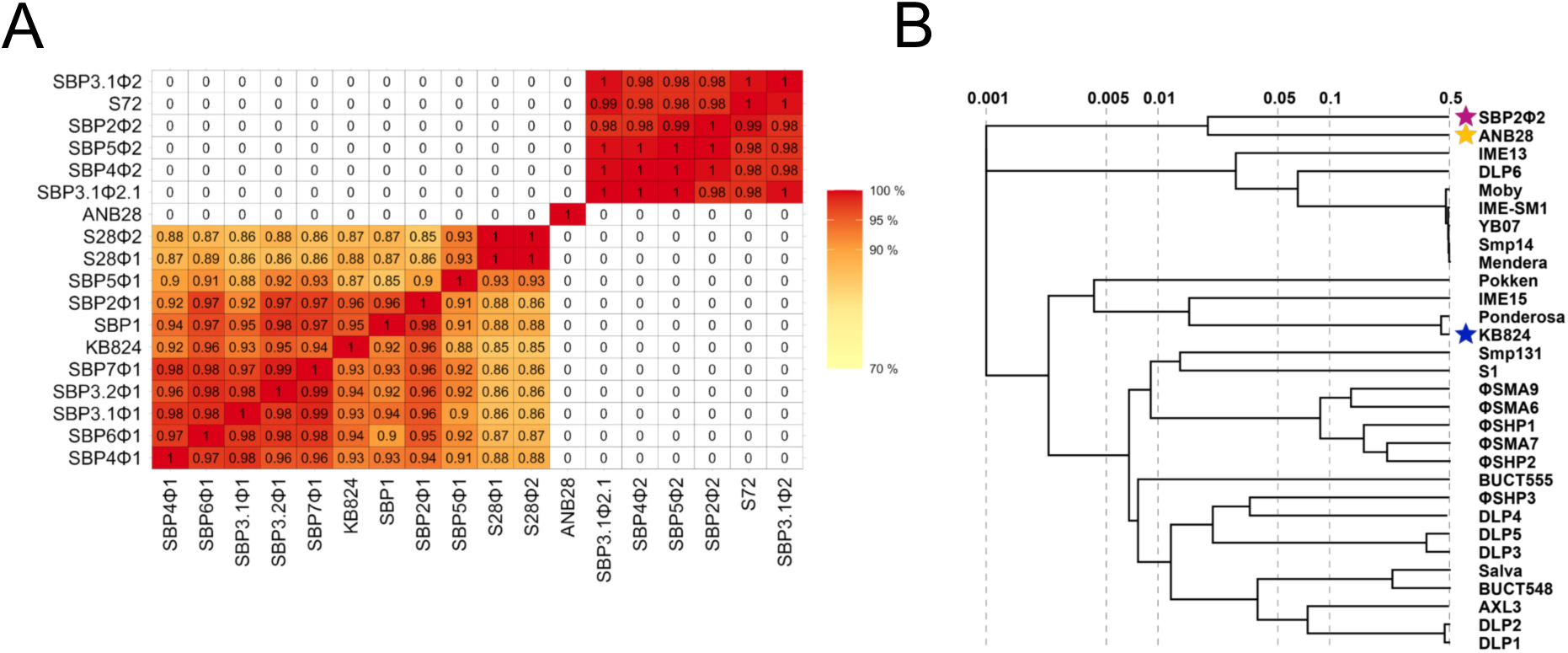
Comparative genomic analysis of in-house STM phages. (**A**) Average Nucleotide Identity Percentage (ANI%) of our 18 isolated STM phages. Phage sequences were cleaned and deduplicated with bbtools^47^, assembled with unicycler^49^, and annotated with RASTtk^51^ to obtain GenBank files. GenBank files were then processed with ANVIO using the <anvi-compute-genome-similarity= option to determine the ANI%^58^. The bottom left cluster represents podoviruses, the top right cluster represents siphoviruses, and the singleton phage is ANB28. (**B**) Phylogenetic Analysis of our three novel STM phages from our own collection against known STM phages previously isolated from the literature^33–41^. Fasta files from previously identified STM phages were obtained from NCBI and were used to generate a phylogenetic tree using VIPtree against our isolated STM phages^43^. (Magenta star indicates SBP2ɸ2; gold indicates ANB28; and blue indicates KB824).

**Figure 2.**
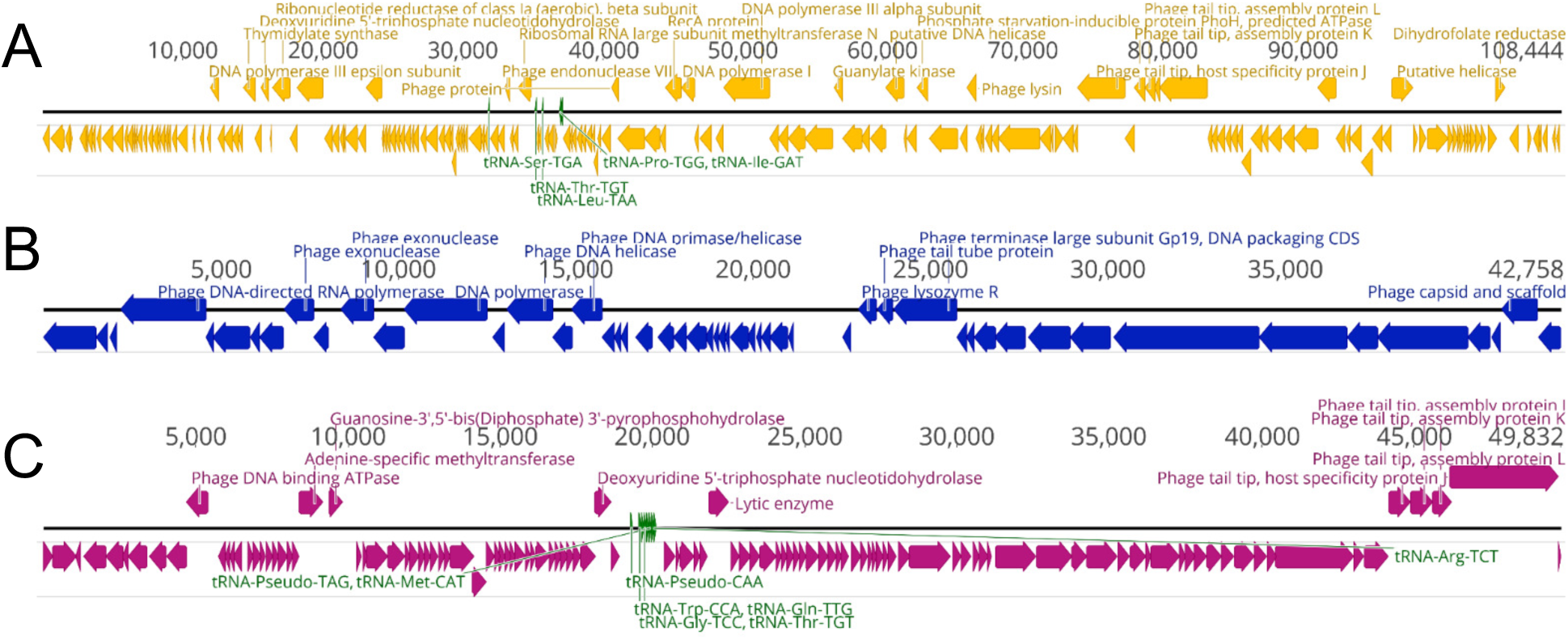
Genomic map of our three unique phages. (**A**) ANB28 (yellow), (**B**) KB824 (blue), and (**C**) SBP2ɸ2 (magenta). GenBank files were visualized in Geneious to establish phage genome maps^59^. The annotated open reading frames (ORFs) are indicated with arrows above or on the black genomic line, while unlabeled hypothetical proteins are shown below. tRNAs are highlighted in green text.

**Figure 3.**
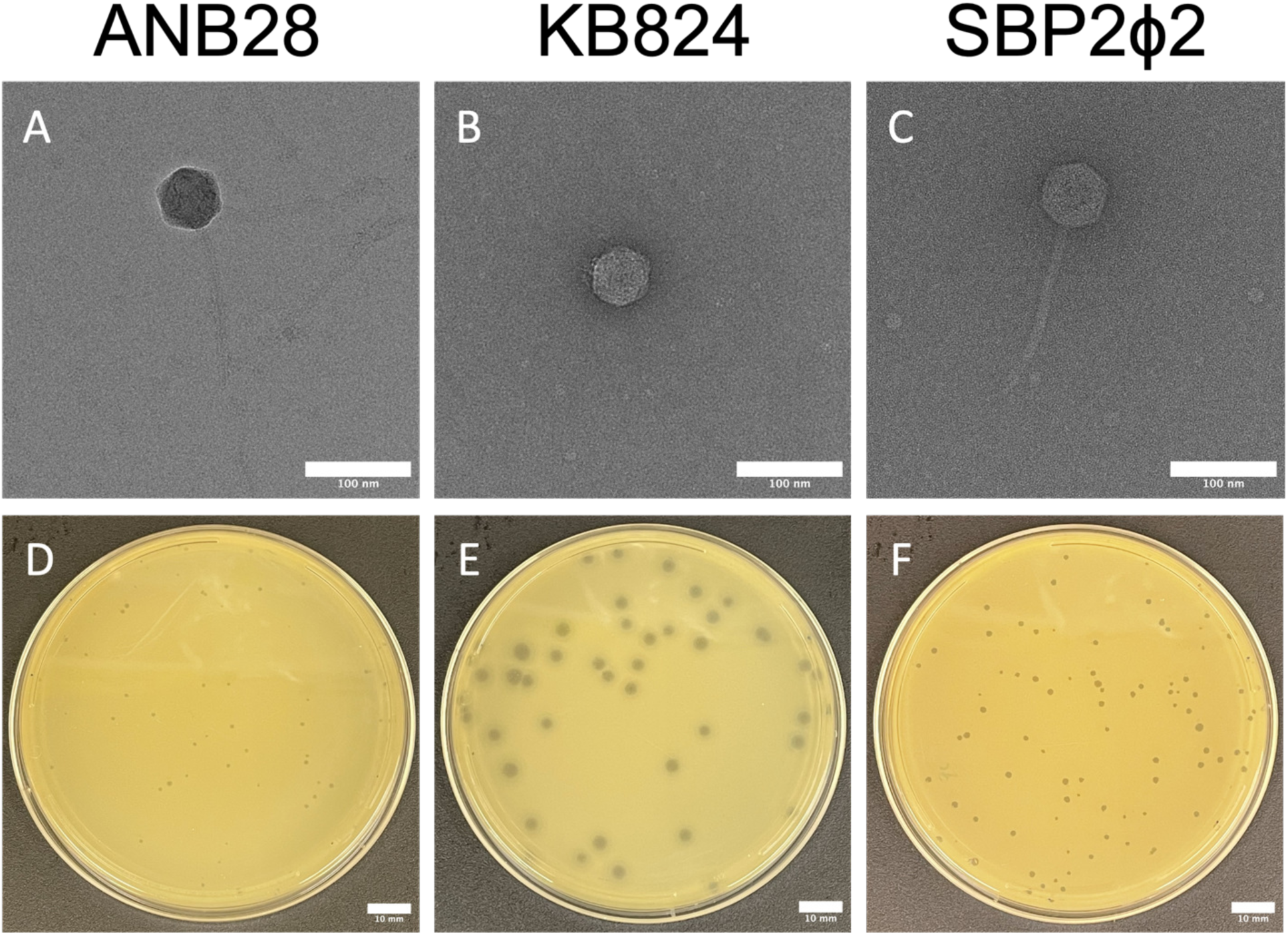
Phage morphology. **(A-C)** Electron Microscopy (EM) micrographs of negatively-stained high-titer phage lysates (>10^8^ PFU/mL) samples. **(D-F)** Phage plaque morphology on Brain Heart Infusion (BHI) soft agar overlay with STM bacterial lawns. Log phase bacteria were mixed with phages at a dilution to achieve countable plaque-forming units (PFUs) and incubated at 37°C for 18-20 hours. Scale bars represent 10 mm. (**A and D**) ANB28 has siphovirus morphology and pinpoint plaques. (**B and E**) KB824 has podovirus morphology and hazy-halo plaques. (**C and F**) SBP2ɸ2 has siphovirus morphology and pleomorphic plaques.

**Table 1.** Summary of Genomic Information. After DNA extraction and assembly, genomes were analyzed for bacterial AMR genes using the Comprehensive Antibiotic Resistance Database (CARD)^54^ and bacterial toxin genes using TAfinder^55^. RASTtk^51^ annotations were performed and assessed in Geneious^59^ for integrase genes, hypothetical proteins (HP), transfer RNAs (tRNAs), open reading frames (ORFs), GC%, and length.

| **Table 1. Genomic Information of Selected STM Phages** |  |  |  |
| --- | --- | --- | --- |
|  | *ANB28* | *SBP2φ2* | *KB824* |
| Length | 108444 | 49832 | 42910 |
| GC% | 53.25 | 52.06 | 59.87 |
| ORFs | 194 | 123 | 76 |
| tRNAs | 5 | 8 | 0 |
| HP | 178 | 114 | 50 |
| Integrase genes | 0 | 0 | 0 |
| AMR genes | 0 | 0 | 0 |
| Toxin genes | 0 | 0 | 0 |
| *HP = Hypothetical Protein, ORF = Open Reading Frame* |  |  |  |

### Basic morphological characterization of three unique STM phages

EM micrographs illustrate ANB28 as having a siphovirus morphology. KB284 and SBP2ɸ2, initially classified based on sequence similarities, were confirmed by EM as having podovirus and siphovirus morphology, respectively. All three phages showed an icosahedral capsid, while both siphoviruses, ANB28 and SBP2ɸ2, contained long, non-contracted tails. KB824, a podovirus, contained a very short non-contracted tail (**Figure 3A-C**). Plaque morphology for each phage was distinct: ANB28 makes pinpoint plaques, KB824 consists of hazy mid-size plaques, and SBP2ɸ2 plaques are clear and pleomorphic (**Figure 3D-F**). KB824 exhibited robust lytic activity at room temperature, showing variation in plaque morphology from a physiologically relevant temperature of 37°C (**Supp Figure 4**).

**Figure 4.**
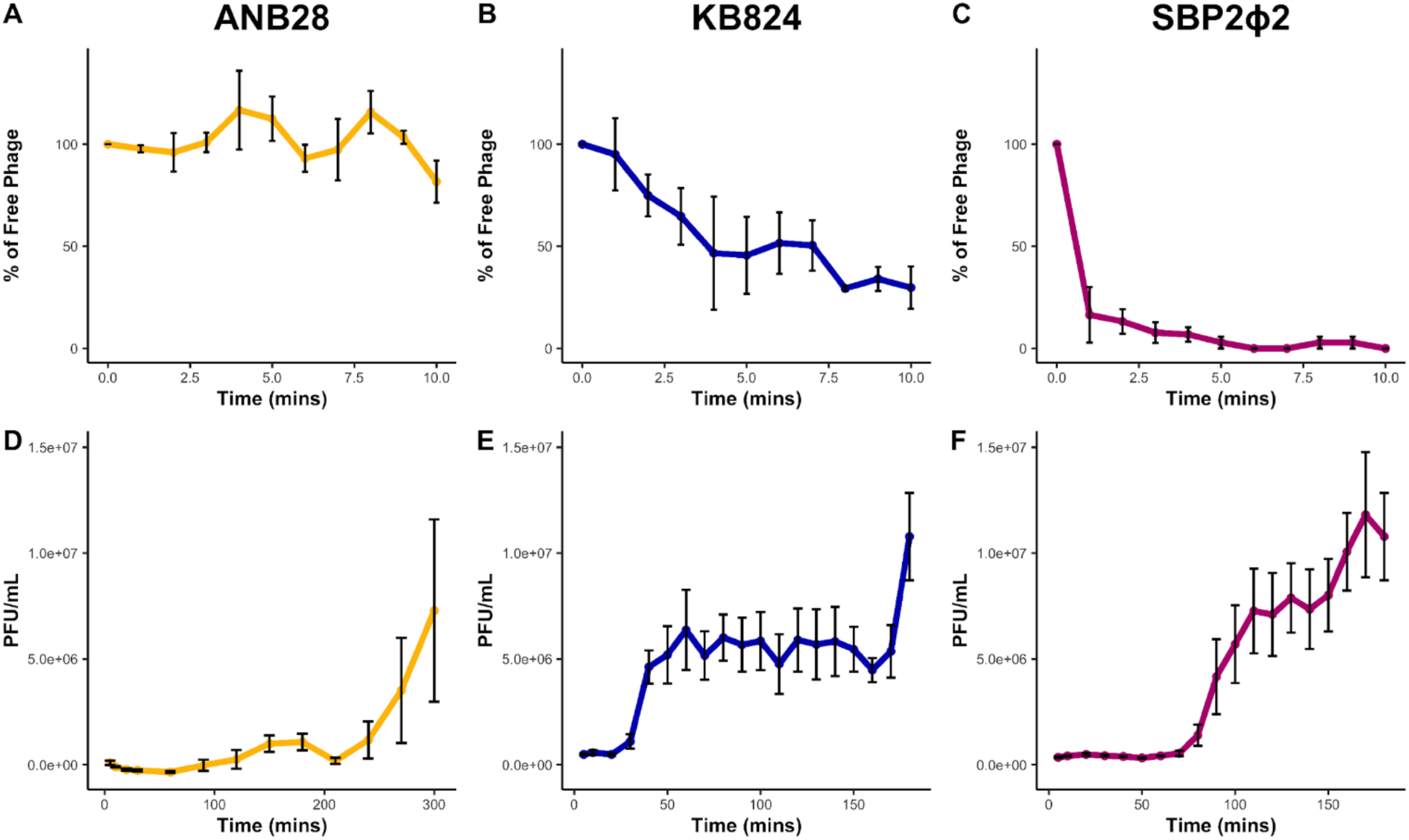
Phage kinetics of STM phages against STM host strain B28B. (**A and D**) ANB28, (**B and E**) KB824, and (**C and F**) SBP2ɸ2. (**A-C**) The rate of phage attachment to bacterial host cells. Sampling was conducted every minute over 10 minutes using log phase bacteria (OD600 0.3) exposed to phage at MOI 0.001^16^. The phage-bacteria mixture samples were first treated with chilled chloroform before they were processed into a soft agar overlay. The percentage of free phage was calculated by dividing the raw data at each time point by the average of the control samples lacking bacteria, multiplied by 100. (**D-F**) One-step growth curves were conducted for three or five hours at 37°C, shaking; samples were taken every 10-30 minutes^17^. Log phase bacteria (OD600 0.3) were exposed to phage at MOI 0.001, and sampling was quickly followed by plating with a soft agar overlay. All plates were incubated at 37°C for 18-20 hours before counting plaques. Three biological replicates were averaged and graphed; error bars represent standard error.

The efficiency of plating (EOP) was conducted against 13 clinically relevant STM strains to assess the host range of each phage in a solid condition using soft agar overlays. A high titer of ANB28 (>10^6^ PFU/mL) was able to infect three STM strains, including the STM-type strain, K279a. KB824 had the broadest host range, with five STM strains susceptible to a 10^5^ PFU/mL titer. SBP2ɸ2 had the narrowest host range, consisting of only two STM strains at a 10^5^ PFU/mL titer (**Table 2**). These results indicate that the three newly discovered phages could infect six of the 13 strains tested on solid media, and each exhibited unique morphology.

**Table 2.**
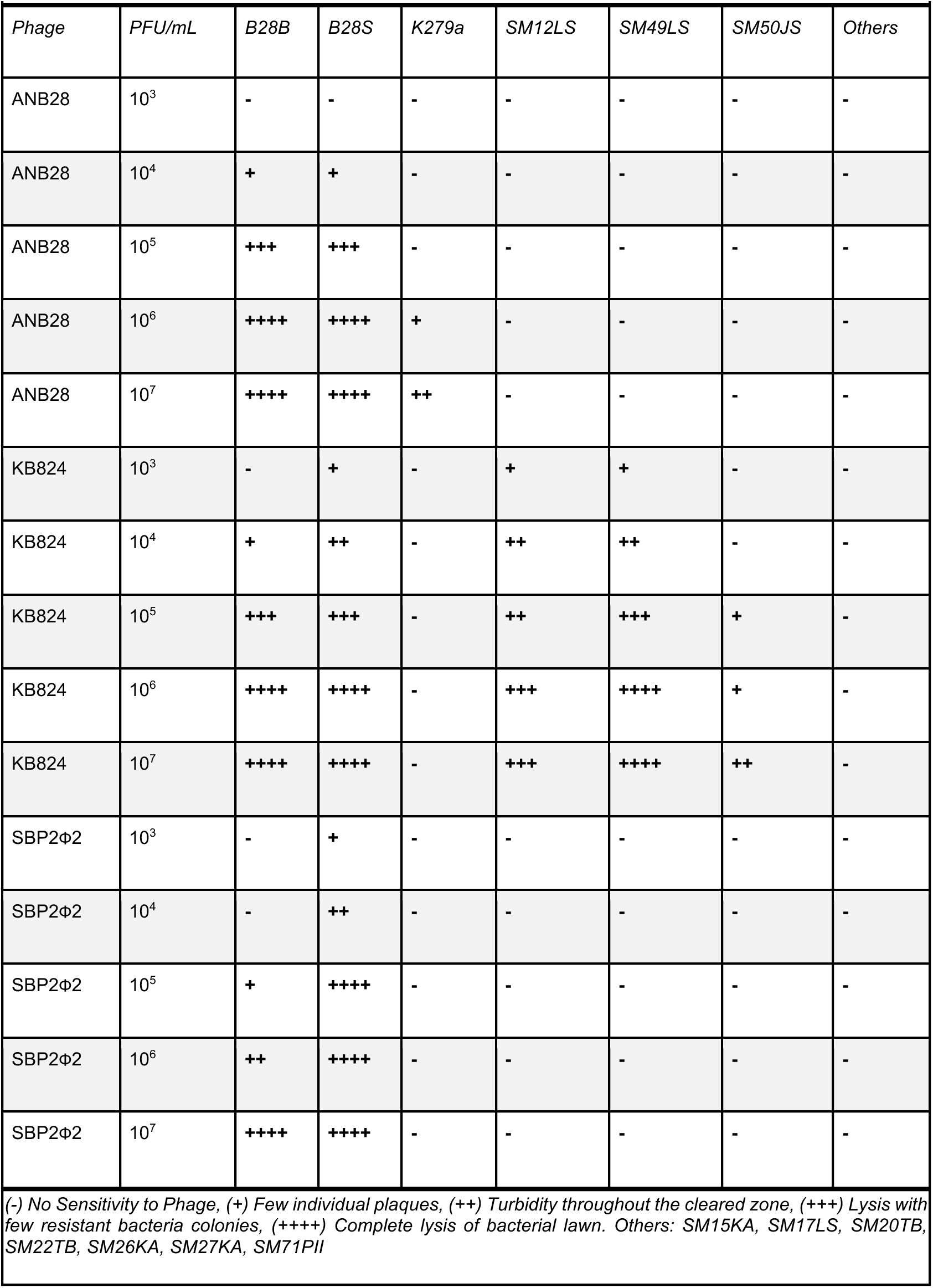
Efficiency of Plating (EOP) for STM Phages on *S. maltophilia* clinical strains.

| **Table 2. Efficiency of Plating for STM Phages on *S. maltophilia* clinical stains** |  |  |  |  |  |  |  |  |
| --- | --- | --- | --- | --- | --- | --- | --- | --- |
| *Phage* | *PFU/mL* | *B28B* | *B28S* | *K279a* | *SM12LS* | *SM49LS* | *SM50JS* | *Others* |
| ANB28 | 10<sup>3</sup> | - | - | - | - | - | - | - |
| ANB28 | 10<sup>4</sup> | + | + | - | - | - | - | - |
| ANB28 | 10<sup>5</sup> | +++ | +++ | - | - | - | - | - |
| ANB28 | 10<sup>6</sup> | ++++ | ++++ | + | - | - | - | - |
| ANB28 | 10<sup>7</sup> | ++++ | ++++ | ++ | - | - | - | - |
| KB824 | 10<sup>3</sup> | - | + | - | + | + | - | - |
| KB824 | 10<sup>4</sup> | + | ++ | - | ++ | ++ | - | - |
| KB824 | 10<sup>5</sup> | +++ | +++ | - | ++ | +++ | + | - |
| KB824 | 10<sup>6</sup> | ++++ | ++++ | - | +++ | ++++ | + | - |
| KB824 | 10<sup>7</sup> | ++++ | ++++ | - | +++ | ++++ | ++ | - |
| SBP2Φ2 | 10<sup>3</sup> | - | + | - | - | - | - | - |
| SBP2Φ2 | 10<sup>4</sup> | - | ++ | - | - | - | - | - |
| SBP2Φ2 | 10<sup>5</sup> | + | ++++ | - | - | - | - | - |
| SBP2Φ2 | 10<sup>6</sup> | ++ | ++++ | - | - | - | - | - |
| SBP2Φ2 | 10<sup>7</sup> | ++++ | ++++ | - | - | - | - | - |
| (-) No Sensitivity to Phage, (+) Few individual plaques, (++) Turbidity throughout the cleared zone, (+++) Lysis with few resistant bacteria colonies, (++++) Complete lysis of bacterial lawn. Others: SM15KA, SM17LS, SM20TB, SM22TB, SM26KA, SM27KA, SM71PII |  |  |  |  |  |  |  |  |

### Infection dynamics of three unique STM phages

Phage kinetic assays, including the rate of attachment and one-step growth curve, were conducted for each of the three STM phages on host bacteria B28B at physiologically relevant body temperatures (∼37°C) using a multiplicity of infection (MOI) of 0.001. The results indicated that each phage attached to host cells at differing rates: SBP2ɸ2 attaching within <5 minutes, KB824 attaching within >10 minutes, and ANB28 demonstrating inefficient attachment to host cells over 10 minutes. KB824 and SBP2ɸ2 both followed the first order of kinetics, while ANB28 showed a slower absorbing subfraction of virions (**Figure 4A-C**). Regarding the one-step growth curve, ANB28 had the most prolonged latent period of around ∼90 minutes, with an average absolute burst size of ∼1x10^6^ PFU/mL for the initial burst. Interestingly, ANB28 returned to a latent phase immediately after the initial burst, followed by a larger burst of progeny virus from the host cell, demonstrating a variable multi-cycle curve. KB824 had the shortest latent period, ∼30 minutes, with an average absolute burst size of ∼5x10^6^ PFUs/mL. SBP2ɸ2 had a latent period of ∼80 minutes with the largest absolute burst size of ∼7x10^6^ PFUs/mL (**Figure 4D-F**). The results indicate that each phage has a unique lytic life cycle regarding the attachment rate, latent period, burst timing, and absolute burst size.

Growth curve analysis of each phage at MOIs of 0.001, 1, and 10 demonstrated that differences in the number of infecting virions for both KB824 and SBP2ɸ2 did not significantly alter the dynamics of infecting host bacteria B28B, as measured in the area under the curve (AUC). KB824 delayed bacterial growth for 10 hours in all MOI conditions (**Figure 5B&E)**. SBP2ɸ2 suppressed bacterial growth for 18-20 hours, with the two higher MOIs matching the exact growth pattern and the lower MOI trending with less reduction in initial growth and delayed bacterial resistance, but no significant differences were identified when AUC was evaluated (**Figure 5C&F**). For ANB28, we observed that MOI 10 caused a significant reduction in overall bacterial growth as measured with the AUC. Surprisingly, for phage ANB28, MOI 0.001 trended longer in preventing resistant bacterial growth than MOI 1; however, there was no significant difference between the two MOIs as measured with AUC (**Figure 5A&D**). These results indicate that, under the tested conditions, the abundance of the three phages has little to no impact on phage predation and phage resistance of host bacteria B28B, as similar growth patterns emerge at the different MOI inputs.

**Figure 5.**
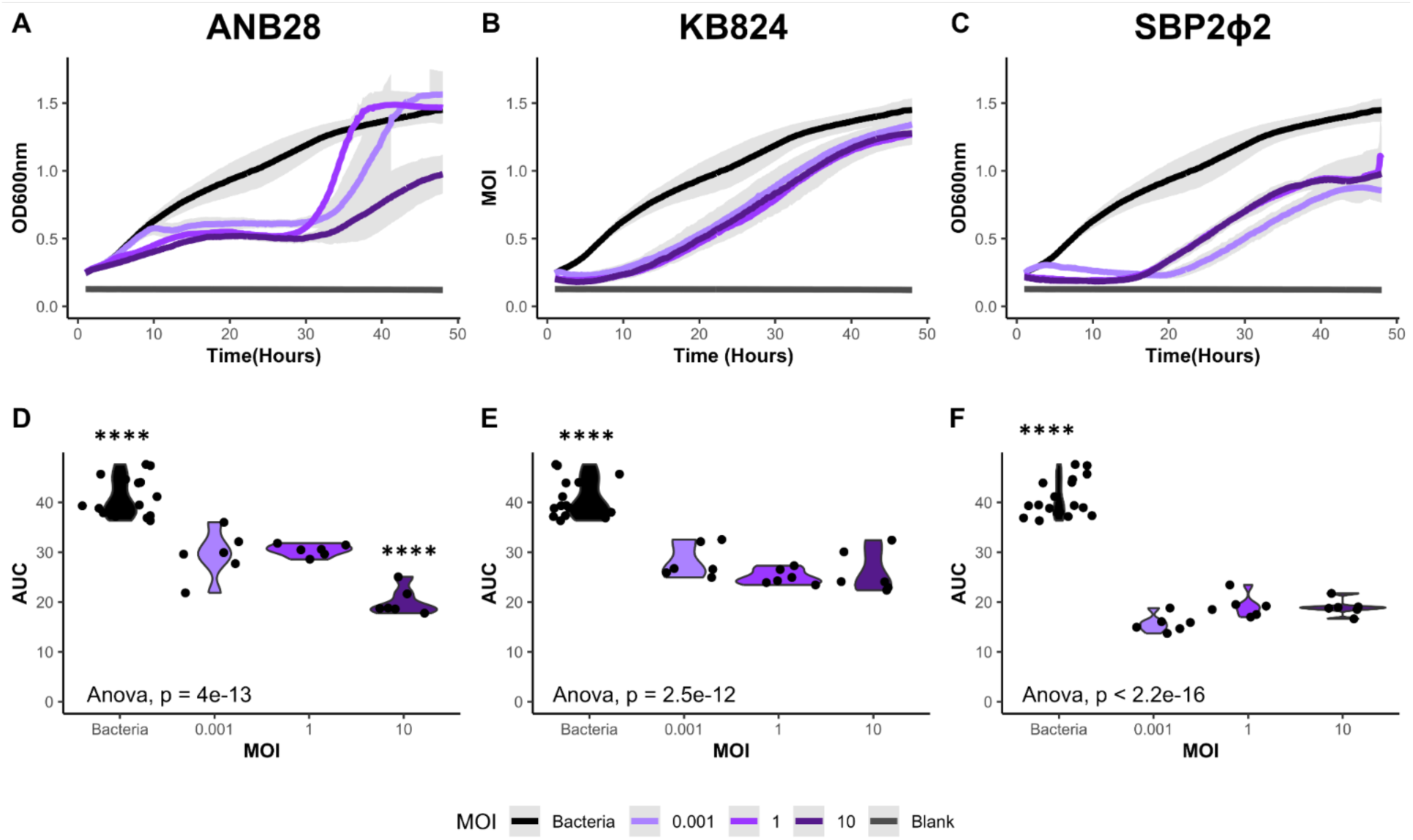
Impacts of varying multiplicity of infection (MOI) against STM host strain B28B. (**A and D**) ANB28, (**B and E**) KB824, and (**C and F**) SBP2ɸ2. (**A-C**) Growth curve analysis (GCA) of bacteria B28B in the presence of three isolated phages. Log phase bacteria (OD600 0.1) were added to 96-well plates and exposed to each phage at three MOIs: 0.001, 1, and 10. Optical density (OD600) was collected in the Agilent LogPhase600 plate reader for 48 hours at 37°C. Averages of the growth curve are graphed with the gray area representing the standard deviation. (**D-F**) A one-way ANOVA was performed using the area under the curve (AUC), calculated with the Growthcurver package in R, after blank adjustment^57,65^. For ANB28, the main effect of MOI is statistically significant and large (F(3, 32) = 58.90, p < .001; Eta2 = 0.85, 95% CI [0.76, 1.00]). Tukey9s HSD Test for multiple comparisons found that the bacterial control and an MOI 10 significantly differed from all other conditions. For SBP2ɸ2, the main effect of MOI is statistically significant and large (F(3, 32) = 172.45, p < .001; Eta2 =0.94, 95% CI [0.91, 1.00]). For KB824, the main effect of MOI is statistically significant and large (F(3, 32) = 51.31, p < .001; Eta2 = 0.83, 95% CI [0.73, 1.00]). For both KB824 and SBP2ɸ2, Tukey9s HSD Test for multiple comparisons found that only the bacterial control significantly differed from all other conditions. Violin plots of AUC are shown with individual data points marked as dots. Data is represented by six growth curves: three biological replicates consisting of two technical replicates each, with bacterial controls assessed on each plate. Light purple represents a low MOI (0.001), purple represents a mid-MOI (1), dark purple represents a high MOI (10), and bacterial control is represented by black. *Significant level*: p < 0.0001(****).

### Infection dynamics of a cocktail comprising three phages with unique genomes and infection kinetics

Growth curve analysis using our three distinct phages combined into a cocktail and the individual phage counterparts was conducted against host bacteria B28B with a combined total MOI of 1. The results indicated that the three-phage cocktail, compared to individual phages, was optimal at suppressing host bacterial growth for an extended period (48 hours) and reducing bacterial resistance in the host bacteria (**Figure 6A**). The AUC of the bacteria-only control was significantly elevated compared to all other conditions. At the same time, the AUC of the three-phage cocktail was significantly decreased compared to all other conditions. The AUC for individual phages varied in significance, with ANB28 and SBP2ɸ2 showing the largest difference in AUC, followed by the AUC for KB824 and SBP2ɸ2, then the AUC for ANB28 and KB824 (**Figure 6B**).

**Figure 6.**
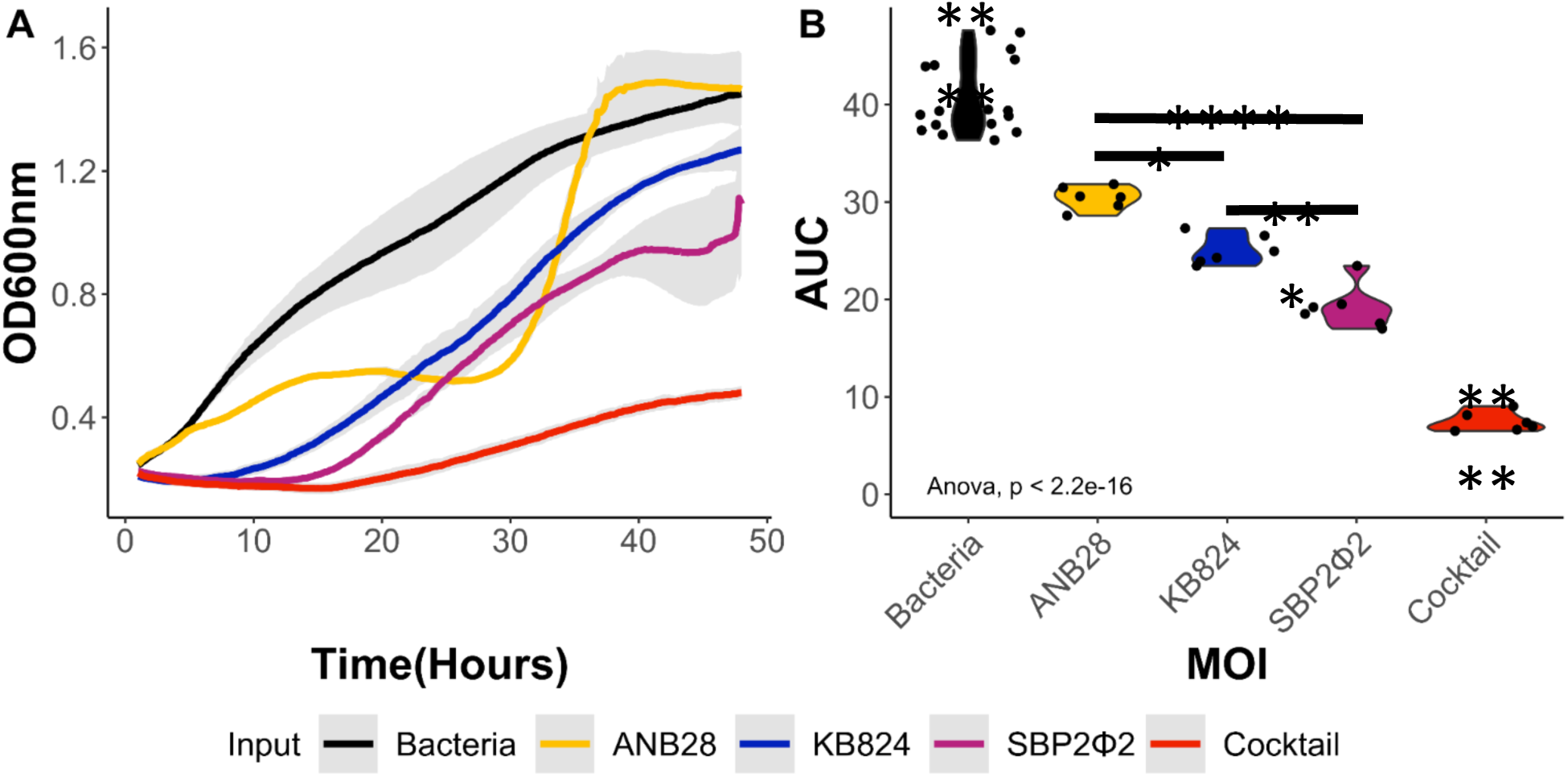
Impact of a three-phage cocktail against STM host strain B28B. (**A**) Growth curve analysis (GCA) of bacteria B28B against a three-phage cocktail consisting of phages ANB28, KB824, and SBP2ɸ2. Log phase bacteria (OD600 0.1) were added to 96 well plates and exposed to individual phages and a three-phage cocktail at an MOI 1. Optical density (OD600) was collected with the Agilent LogPhase600 plate reader for 48 hours at 37°C. Averages of the growth curves are graphed with the gray area representing the standard deviation. (**B**) A one-way ANOVA was performed using the area under the curve (AUC) after blank adjustment, calculated using the Growthcurver package in R^57,65^. The main effect of Input is statistically significant and large (F(4, 37) = 191.63, p < .001; Eta2 = 0.95, 95% CI [0.93, 1.00]). Tukey9s HSD Test for multiple comparisons found that all conditions were statistically different, with the bacterial control and cocktail conditions having the most significant difference. Violin plots of AUC are shown with individual data points marked as dots. Data is an average of six growth curves comprising two technical replicates across three biological runs with bacterial controls assessed on each plate. Bacteria (black), ANB28 (yellow), KB824 (blue), SBP2ɸ2 (magenta), and the three phage cocktail (red). *Significant levels*: p<0.05 (*), p < 0.001 (***) and p < 0.0001(****).

Extensive host range analysis was performed with the three-phage cocktail and individual phages against 46 clinically relevant STM strains at 37°C in liquid culture at an MOI 1 (based on host bacteria B28B) (**Supp Fig. 5**) AUC was calculated for 12, 20, and 40 hours before blank adjustment. The growth percentage was normalized to the bacteria-only condition to evaluate lytic activity in a strain-dependent manner using the following equation: [(1-(AUC_control_ - AUC_phage_)/AUC_control_)*100]. The reduction in the red opacity indicates a reduction in bacterial growth; thus, lighter shades of red represent an increase in lytic phage activity. Approximately half of the evaluated STM strains succumb to phage infection under cocktail conditions (**Supp Fig. 5**). These results suggest phage infectivity is highly selective; however, we see reduced phage resistance and bacterial growth when multiple phages can infect a bacterial strain. Data from six strains in which the three-phage cocktail showed a reduction in bacterial growth at the 40-hour time period, compared to individual phages, were further analyzed with growth curve analysis (**Figure 7**). The three-phage cocktail prevented the development of phage-resistance altogether, except for SM16LS, which caused a large delay in bacterial growth. These results highlight the enhanced efficiency of a multi-phage cocktail against bacterial suppression, indicating a potential strategy for mitigating phage resistance.

**Figure 7.**
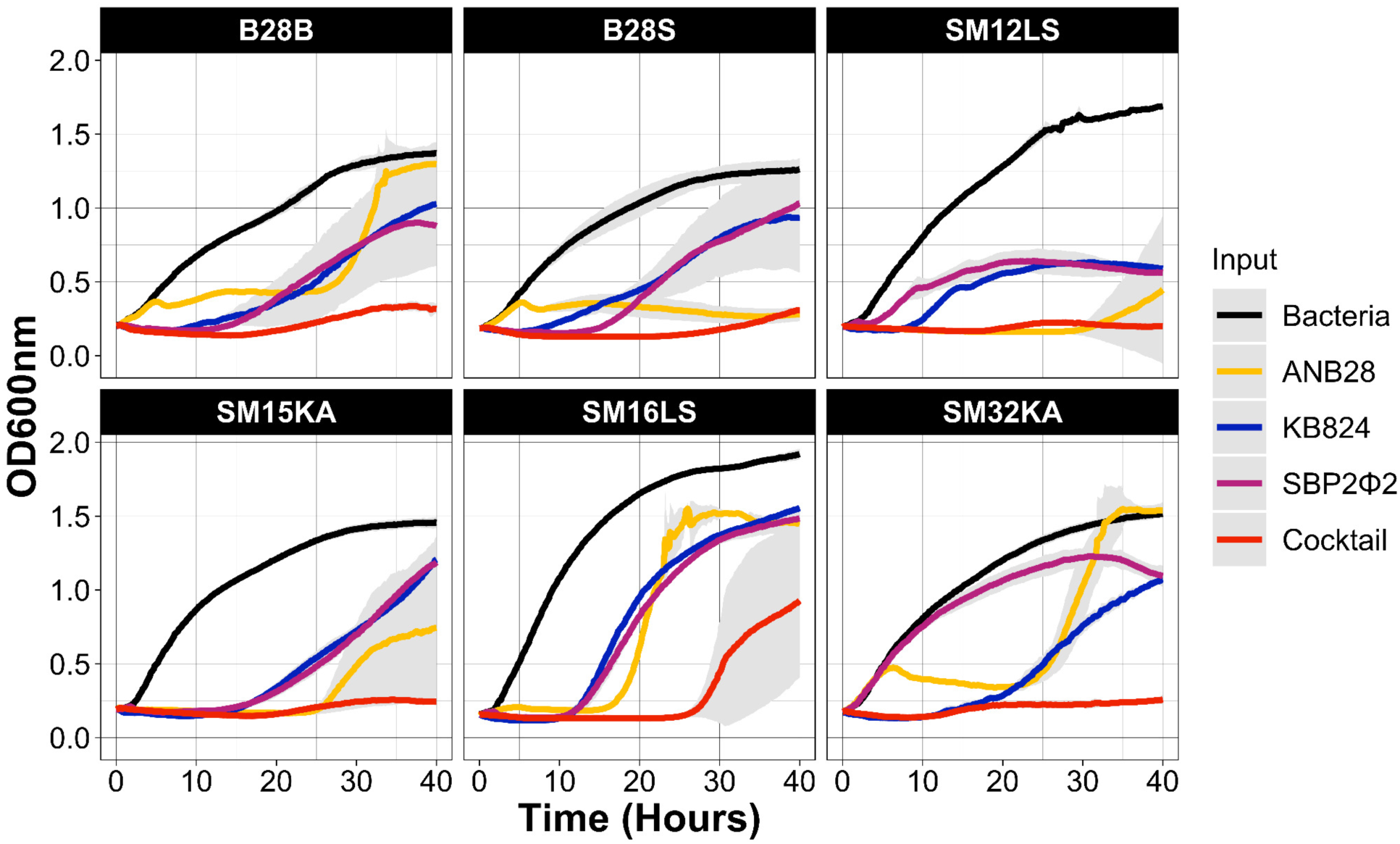
The impacts of a three-phage cocktail against six clinically relevant STM strains. Extensive host range analysis of our STM phages, both individually and in three-phage cocktail, against 46 STM strains are illustrated in Supp Fig 5. STM strains were grown to a log phase (OD600 0.1) and exposed to ANB28, KB824, SBP2ɸ2, and a three-phage cocktail comprising all three phages in a 96-well plate setup. Phages were exposed at MOI 1 based on the titer of host strain B28B. Growth curve data (OD600) was collected from the Log Phase600 plate reader for 40 hours at 37°C. The results are presented as the average of three technical replicates, with the standard deviation denoted in gray. Graphical analysis was conducted in R using ggplot2.

## DISCUSSION

Here, we analyzed a collection of STM phages; three genetically distinct clusters were identified out of 18 initially harvested phage isolates, with ANB28 being genetically unique. Based on the genetic analysis, three representative phages —ANB28, KB824, and SBP2ɸ2— were selected for further evaluation. Phylogenetic analysis confirmed KB824 was closely related to *Stenotrophomonas* phage Ponderosa, while ANB28 and SBP2ɸ2 diverged from previously isolated STM phages. All three phages were free of genome-encoded integrase, AMR, or toxin genes. Phenotypic observations demonstrated distinct plaque morphology for each phage, while EM confirmed phage morphologies as podovirus for KB824 and siphoviruses for ANB28 and SBP2ɸ2. Phage kinetic revealed each phage had a unique attachment rate and life cycle when targeting host bacteria, B28B; however, when combined into a three-phage cocktail, the phages significantly reduced B28B growth and effectively mitigated phage resistance. While the host range analysis revealed a unique and narrow profile for each phage, their collective efficacy exhibited a notable reduction in phage resistance when the bacterial strain was susceptible to multiple phages, which was highlighted in six clinically relevant STM strains.

Establishing safety guidelines for phage for therapeutic use is an important ongoing effort^11^. We have screened our phages to the best of our ability to ensure their safety, with no identifiable toxin, AMR, or lysogenic lifestyle-associated genes. The three representative phages we chose are distinct in genetic makeup, phenotypic observations, infection kinetics, and host range, highlighting the spectrum of phage diversity and the mechanisms each phage operates. Interestingly, ANB28 exhibited a higher degree of uniqueness as a singleton phage within its cluster, and BLASTn analysis revealed low similarities to any known phage. While it may feel surprising to find a novel phage from urban wastewater, phage diversity remains unexplored, and this is consistent with the inherent diversity of phages^7^. Additionally, the impacts of a multi-phage cocktail on a susceptible host bacteria provide supporting evidence that genetically distinct phages give rise to unique infection kinetics, facilitating lytic activity to overpower bacterial growth and resistance compared to an individual phage. This supports our initial hypothesis. Differences in phage attachment and host range highlight the complexity of phage-bacteria interactions. However, these observations, specifically the phage host range against 46 clinically relevant STM strains, could be attributed to bacterial host factors, not the phage, such as genetic mutations or adaptation associated with phage defense systems. These host elements could play a major role in which bacteria can be infected by which phage. Thus, further investigation into bacterial host defenses is warranted to understand how we can optimize phage applications.

Identifying genetically distinct phages, characterizing their infection parameters, and evaluating the efficacy of a multi-phage cocktail demonstrate promising strategies for optimizing phage-based applications. However, other strategies that have proven to be successful with antibiotics may also be adapted for phage treatment, such as cycling and switching treatment approaches as resistance emerges^44^. Although this approach has the potential to increase bacterial killing, the strategy is complex and requires real-time data analysis for isolated bacterial cultures, which could delay treatment. Therapeutic failure in antibiotic treatment of recalcitrant STM infections renders limited options for patients; however, phages could become a critical, life-saving strategy. Our research reaffirms the importance of precision medicine in phage therapy, demonstrating the potential benefits of tailoring phage cocktails to specific bacterial strains, thereby enhancing treatment efficiency and mitigating the development of phage resistance^45^. Additionally, screening phages devoid of AMR genes is essential to establishing phage therapy as a practical solution that would provide a vital foundation for evaluating preclinical safety, efficacy, and feasibility. Establishing practical phage applications could have extensive implications for public health and mortality rates and reduce healthcare costs associated with recurring AMR infections. Additionally, this research provides insight into phage-bacteria interactions, highlighting the critical time points pertinent to the phage replication strategies and laying the groundwork for future studies. By advancing our knowledge of phage-bacteria interactions, we hope to provide insights into phage biology and potential strategies for optimization phage applications.

Our research highlights the importance of susceptibility testing prior to phage therapy to ensure a phage will be successfully matched for bacterial clearance. Additionally, further development of our STM phage library is critical to comprehensively cover the diversity of our STM strain collection. Thus, a limitation of our research is the dearth of STM phages, which may only provide a glimpse of phage diversity in the host STM. However, we conducted phylogenetic analysis with 27 previously identified STM phages to understand STM phage diversity. Additionally, this research consisted of laboratory-based experiments, which do not directly translate into real-world clinical settings. Therefore, further exploration and validation is necessary to confirm the applicability of our findings. However, screening phage information before the clinical trial is necessary and cost-effective for establishing foundational research. Lastly, we were able to show the impacts of a three-phage cocktail; however, we must investigate the specific interactions within the cocktail. Understanding the synergistic effects of different phages within a cocktail is vital in optimizing therapeutic applications. Our future studies will explore this phage complexity using host bacteria B28B gene expression profiles under individual and cocktail phage predation.

Our research attempts to understand phage cocktail dynamics in a poorly studied opportunistic pathogen, STM. Through our study, we were able to demonstrate (1) successful screening and selection of STM phages, (2) identification of phage diversity in terms of genomics and kinetics, and (3) establishment of an effective phage cocktail against host bacteria B28B. This data warrants future research into phage-bacteria mechanism, evolved phage-resistance, and phage-delivery methods. Future studies will involve a transcriptomic analysis of host bacteria B28B under phage predation in a time-dependent manner, both with an individual phage and in a cocktail setting, which will aid in understanding the replication strategy of the phage and the potential vulnerabilities of the host bacteria. Additionally, investigation into delivery methods will be essential as this will provide insight into phage stability and effectiveness, which could be vital in targeting infection burdens.

## MATERIAL AND METHODS

### Bacteria Cultures

The bacterial strains used in this study are listed in Supplemental Table S1. STM strain B28B was isolated from an oncology patient at UCSD (Summer, 2020). B28B was grown in Brain Heart Infusion Broth (BHI; Research Products International) at 37°C on a 200-rpm shaker. Glycerol stocks were made at a final glycerol concentration of 25%. Bacteria were grown by streaking from glycerol stocks onto BHI plates and incubated at 37°C for 18-20 hours. For experiments and assays, isolated colonies were grown in overnight broth culture; the next day, a 1:10 or 1:20 dilution into BHI was placed at 37°C on a 200 rpm shaker to achieve a log phase at OD600 of 0.3 or 0.1, respectively.

### Phage Lysate, Titering, and Plaque Morphology

The phage isolates used in this study are listed in Supplemental Table S2. Phage propagation was based on Bonilla et al., 2016.^46^ Phage lysates were stored with a final concentration of 10% glycerol at -80°C. Phage titering was done every two weeks and recorded over time based on when the phage was harvested, which correlated to a specific lot number. For plaque morphology and phage titering, serial dilutions of phage lysates were used to achieve a countable plaque number. Plating consisted of 10 uL of the diluted lysate against 100 uL of log phase host bacteria B28B using a BHI soft agar overlay incubated at 37°C for 18-20 hours. Three technical replicates were averaged to calculate the PFU/mL of the stock concentration of a lysate or scanned for plaque morphology. The phage titer of KB824 for temperature assessment was conducted similarly. Duplicate plates were made; one set was incubated at 37°C while the other set was incubated at room temperature for 18-20 hours, then scanned. Scanned was performed using the EPSON Perfection V600 Photo Scanner.

### Sequencing and Bioinformatics

The phages used in this study are listed in Supplemental Table S2. Phage DNA was extracted from high-titer stocks using a QIAamp UltraSens Virus kit (Qiagen, Cat. 53706) per the manufacturer’s instructions. Before performing the DNA extraction, all phages were treated with 2 uL of RNAase A (50,000 U/mL, New England BioLabs, Cat. M02403S) and 50 uL of NEB buffer, followed by 5 uL of DNAase I (2000 U/mL, New England BioLabs, Cat. M0303S). Samples were then treated with 50 uL of NEB buffer for a 30-minute incubation at 37°C followed by a 10-minute incubation at 74°C to inactive the enzymes. The extracted DNA was quantified using a Qubit dsDNA High Sensitivity assay kit (Invitrogen, Cat. Q32851), and library preparation was done using the Nextera XT DNA LP kit (Illumina). Sequencing was performed on Illumina’s iseq100 using a paired-end approach (2*150 bp). Raw Illumina reads were uploaded to the High-Performance Community Computing Cluster (HPC3) and cleaned with “bbduck,” and duplicates were removed with “dedup,” both from bbtool^47^. Human contamination was removed with Bowtie2 v2.4.1^48^. Reads were assembled with unicycler^49^, checked for quality with QUAST^50^, and annotated with RASTtk^51^. Coverage analysis was used to identify the contig of interest if sequencing resulted in multiple contigs. Fasta files were concatenated and uploaded to the VIRIDIC server for the VIRIDIC analysis^52^, while CheckV analysis was run in the command line^53^. Phage therapy candidacy screening of fasta file for AMR genes and toxin-encoding genes was accomplished using the CARD database^54^ and TAfinder^55^, respectively. Phylogenetic analysis was performed in VIPtree^43^ using fasta files from a compiled list of STM phages collected from literature sources^33–41^. Coverage plots were performed by mapping clean reads to a Bowtie2 database for each phage fasta file^48^. Samtools was then used for read counts^56^, while data visualization was done in R^57^. Comparative genomics of GenBank files was accomplished with Anvi’o using the ANI% option (“anvi-compute-genome-similarity”) and visualized using their established interface^58^. Output for ANI% was visualized in R. Genome maps were visualized in Geneious Prime^59^ with GenBank files, and manual checks were performed for integrase genes, hypothetical proteins (HP), transfer RNAs (tRNAs), open reading frames (ORFs), GC%, and genome length.

### The Efficiency of Plating

B28B was grown to log phase (OD600 0.3) in BHI broth. Molten agar overlays of 4.9 mL were performed on square petri plates (VWR Cat. 60872-310) using 140 uL of bacteria culture and allowed to solidify at room temperature for 40 minutes. Phage stocks were processed to a 10^7^ PFU/mL titer, and serial dilutions were made to achieve 1 x 10^6^, 1 x 10^5^, 1 x 10^4^, and 1 x 10^3^ PFU/mL titer. Aliquots of the phage dilution were added to the bacterial lawn in a 3 µL volume and allowed to dry. Plates were incubated at 37°C for 18-20 hours before being scored for lysis based on a published protocol^60^. Each phage dilution was run in technical duplicate against 13 different clinically relevant STM strains.

### Phage Morphology by Electron Microscopy

After phage propagation, to observe virion morphology, samples were negatively stained using established procedures^61^ (briefly summarized here) and imaged by Electron Microscopy. 200-mesh Gilder copper grids (Ted Pella) were carbon-coated in-house, and 0.75% Uranyl Formate stain was prepared fresh. Grids were negatively glow-discharged using a PELCO easiGlow (Ted Pella) prior to staining. Samples were stained as-is and by using a dilution series to avoid potential overpacking. 3 μL of each sample was applied to a grid and allowed to adsorb for 10 seconds before excess liquid was removed using filter paper, washed twice with Milli-Q water, stained using 0.75% Uranyl Formate, and allowed to air dry. All grids were imaged, and data was collected using a JEOL JEM-2100F transmission electron microscope equipped with a Gatan OneView 4k x 4k camera. Scale bars in Figure 3 A-C were added using ImageJ^62^.

### Rate of Attachment

The rate of attachment was based on Kropinski et al., 2009, with minor modifications^16^. B28B was grown to a log phase (OD600 0.3) in BHI broth. The absorbance flask (9 mL of bacteria) and media flask (9 mL of BHI) were equilibrated for 5 minutes at 37°C and shaken at 200 rpm (Entech Instruments 5600 SPEU) before 1 x 10^5^ PFU were added to both flasks (t=0). Vials of 50 µL of CHCl_3_ and 950 µL of BHI were chilled for 10 minutes before adding 50 µL of bacteria-phage mixture. Sampling was performed every 10 minutes, vortexed, and placed on ice. Controls were sampled and processed after the 10-minute experimental samples were obtained, as previously described for the experimental conditions. The molten overlay was performed chronologically for each time point and the two controls. Petri plates solidified at room temperature (RT) for 40 minutes and then incubated at 37°C for 18-20 hours. Data of absolute PFUs were recorded and converted into percentages of free phage by dividing the average control value. Each phage isolate was performed against three biological replicates of host bacteria, B28B.

### One-Step Growth Curve

This protocol was performed with minor adjustments based on Kropinski et al., 2018^17^. B28B was grown to log phase (OD600 0.3) in BHI broth. An adsorption flask was prepared with 900 µL of bacteria, while the dilution flasks (10^-2^ flask and 10^-4^ flask) were prepared with 9.9 mL of fresh BHI. All flasks were placed on a shaker (∼200 rpm) to equilibrate to 37°C (Entech Instruments 5600 SPEU). Phage was added to the adsorption flask at an MOI of 0.001 in a 100 µL volume and mixed well. Immediately afterward, 100 µL was taken from the adsorption flask, added to the 10^-2^ flask, and mixed well; this process was repeated from the 10^-2^ flask to the 10^-4^ flask. For phage ANB28, a 10^-3^ flask was prepared. Directly following, 2 mL of the 10^-4^ flask (for phage ANB28, 10^-3^ flask) was removed and added to a microcentrifuge tube containing chilled CHCl_3_. At specific time points, aliquots of either 500 µL, 250 µL, 100 µL, or 50 µL were taken from the diluted flask, which was then used in the molten agar overlay with host bacteria to achieve countable plaques. Upon completion of the phage-bacteria sampling, either 500 µL, 250 µL, 100 µL, or a combination of the two were taken from the CHCl_3_-treated control and processed, as previously stated. Petri plates were allowed to solidify at RT and then incubated at 37°C for 18-20 hours. Absolute PFUs were counted and calculated into PFU/mL with averaged control values of two duplicates subtracted from each data point and then graphed.

### MOI and Cocktail Growth Curves

B28B was grown to log phase (OD600 0.1) in BHI broth. A 96-well plate with a water perimeter (∼200 µL/well) to reduce experiment evaporation was used. Media controls and bacterial aliquots of 180 µL were placed into designated wells. Phage lysates were diluted in SM buffer to achieve an MOI of 0.001, 1, 10 in a 20 µL aliquot. The MOIs were held constant for the three-phage cocktail, incorporating a third of each phage. Either phage dilutions or SM buffer was placed in the designated wells. Plates were run on the Agilent LogPhase600 for 48 hours at 37°C. Data was graphed in R using “dplyr” and “ggplot2” to assess for bacterial contamination in a 96-well plate layout^63,64^. AUC was determined with the Growthcurver package^65^. Statistics were conducted in R^57^ using a one-way ANOVA to determine if the AUC for each phage input differed. A Post Hoc test was performed to identify which conditions and phages were statistically different.

### Host Range Growth Curves

All STM strains were grown to a log phase (OD600 0.1) in BHI broth. Each STM strain was exposed to ANB28, KB824, SBP2Φ2, and a combination of the three phages at an MOI of 1 based on the host strain B28B in technical triplicates. A 96-well plate with water (∼200 µL/well) in the top and bottom rows was used to reduce evaporation. Media controls and bacterial aliquots of 180 µL were placed into designated wells. Phage lysates were diluted in SM buffer to achieve a 20 µL aliquot, and either phage dilutions or SM buffer was placed into the designated wells. An MOI of 1 was held constant for the three-phage cocktail, incorporating a third of each phage. Plates were run on the Agilent LogPhase600 for 48 hours at 37°C. Data was graphed in R using “dplyr” and “ggplot2” to assess for bacterial contamination in a 96-well plate layout^63,64^. AUC was calculated using “gcplyr,” and technical replicates were averaged after removing the blank^66^. Growth percentage was calculated using the following equation: Growth% = (1-(Average Bacteria only AUC - Average Phage AUC)/Average Bacteria only AUC)*100, and data was visualized with heatmaps.

## DATA AVAILABILITY

The code for analyzing and making figures is available at https://github.com/amonsiba/STM_phage_cocktail. Raw sequencing data has been uploaded to the SRA under BioProject PRJNA1121625.

## ACKNOWLEDGEMENTS

We would like to acknowledge the City of Escondido, CA Wastewater Division for providing influent samples from which phages were isolated. Alisha N. Monsibais was supported with a graduate fellowship from the National Institute of Allergy and Infectious Diseases (NIAID; AI141346). Katrine Whiteson and David Pride were funded by an R21 award from NIAID (5R21AI149354-02). Sage Dunham received support through the Cystic Fibrosis Foundation for a Postdoctoral Fellowship award (CFF 003135F221). Diana S. Suder was supported with a graduate fellowship from the Graduate Assistance in Areas of National Need (GAANN; P200A210024) provided by the U.S. Department of Education. The Gonen Lab is supported by the National Institute of General Medical Sciences, grant R35-GM142797.

## REFERENCES

1. Murray, C. J. L. et al. Global burden of bacterial antimicrobial resistance in 2019: a systematic analysis. The Lancet 399, 629–655 (2022).

2. Dadgostar, P. Antimicrobial Resistance: Implications and Costs. Infect. Drug Resist. Volume 12, 3903–3910 (2019).

3. Prestinaci, F., Pezzotti, P. & Pantosti, A. Antimicrobial resistance: a global multifaceted phenomenon. Pathog. Glob. Health 109, 309–318 (2015).

4. Piddock, L. J. The crisis of no new antibiotics—what is the way forward? Lancet Infect. Dis. 12, 249–253 (2012).

5. Ventola, C. L. The antibiotic resistance crisis: part 1: causes and threats. P T Peer-Rev. J. Formul. Manag. 40, 277–283 (2015).

6. Gordillo Altamirano, F. L. & Barr, J. J. Phage Therapy in the Postantibiotic Era. Clin. Microbiol. Rev. 32, e00066–18 (2019).

7. Clokie, M. R., Millard, A. D., Letarov, A. V. & Heaphy, S. Phages in nature. Bacteriophage 1, 31–45 (2011).

8. Young, R., Wang, I.-N. & Roof, W. D. Phages will out: strategies of host cell lysis. Trends Microbiol. 8, 120–128 (2000).

9. Nobrega, F. L. et al. Targeting mechanisms of tailed bacteriophages. Nat. Rev. Microbiol. 16, 760–773 (2018).

10. Summers, W. C. The strange history of phage therapy. Bacteriophage 2, 130–133 (2012).

11. Suh, G. A. et al. Considerations for the Use of Phage Therapy in Clinical Practice. Antimicrob. Agents Chemother. 66, e02071–21 (2022).

12. Maddocks, S. et al. Bacteriophage Therapy of Ventilator-associated Pneumonia and Empyema Caused by *Pseudomonas aeruginosa*. Am. J. Respir. Crit. Care Med. 200, 1179–1181 (2019).

13. Hoyle, N. et al. Phage therapy against Achromobacter xylosoxidans lung infection in a patient with cystic fibrosis: a case report. Res. Microbiol. 169, 540–542 (2018).

14. Saussereau, E. et al. Effectiveness of bacteriophages in the sputum of cystic fibrosis patients. Clin. Microbiol. Infect. 20, O983–O990 (2014).

15. Ferry, T. et al. Personalized bacteriophage therapy to treat pandrug-resistant spinal Pseudomonas aeruginosa infection. Nat. Commun. 13, 4239 (2022).

16. Kropinski, A. M. Measurement of the Rate of Attachment of Bacteriophage to Cells. in Bacteriophages (eds. Clokie, M. R. J. & Kropinski, A. M.) vol. 501 151–155 (Humana Press, Totowa, NJ, 2009).

17. Kropinski, A. M. Practical Advice on the One-Step Growth Curve. in Bacteriophages (eds. Clokie, M. R. J., Kropinski, A. M. & Lavigne, R.) vol. 1681 41–47 (Springer New York, New York, NY, 2018).

18. Hyman, P. Phages for Phage Therapy: Isolation, Characterization, and Host Range Breadth. Pharmaceuticals 12, 35 (2019).

19. Chan, B. K., Abedon, S. T. & Loc-Carrillo, C. Phage cocktails and the future of phage therapy. Future Microbiol. 8, 769–783 (2013).

20. Hampton, H. G., Watson, B. N. J. & Fineran, P. C. The arms race between bacteria and their phage foes. Nature 577, 327–336 (2020).

21. Dupuis, M.-È., Villion, M., Magadán, A. H. & Moineau, S. CRISPR-Cas and restriction– modification systems are compatible and increase phage resistance. Nat. Commun. 4, 2087 (2013).

22. Lopatina, A., Tal, N. & Sorek, R. Abortive Infection: Bacterial Suicide as an Antiviral Immune Strategy. Annu. Rev. Virol. 7, 371–384 (2020).

23. Dy, R. L., Richter, C., Salmond, G. P. C. & Fineran, P. C. Remarkable Mechanisms in Microbes to Resist Phage Infections. Annu. Rev. Virol. 1, 307–331 (2014).

24. Yang, Y. et al. Development of a Bacteriophage Cocktail to Constrain the Emergence of Phage-Resistant Pseudomonas aeruginosa. Front. Microbiol. 11, 327 (2020).

25. Naknaen, A. et al. Combination of genetically diverse Pseudomonas phages enhances the cocktail efficiency against bacteria. Sci. Rep. 13, 8921 (2023).

26. Wandro, S. et al. Phage Cocktails Constrain the Growth of *Enterococcus*. mSystems 7, e00019–22 (2022).

27. Sanchez, B. C. et al. Development of Phage Cocktails to Treat E. coli Catheter-Associated Urinary Tract Infection and Associated Biofilms. Front. Microbiol. 13, 796132 (2022).

28. Brooke, J. S. Stenotrophomonas maltophilia: an Emerging Global Opportunistic Pathogen. Clin. Microbiol. Rev. 25, 2–41 (2012).

29. Gröschel, M. I. et al. The phylogenetic landscape and nosocomial spread of the multidrug-resistant opportunist Stenotrophomonas maltophilia. Nat. Commun. 11, 2044 (2020).

30. Yang, Z. et al. Prevalence and detection of Stenotrophomonas maltophilia carrying metallo-Î^2^-lactamase blaL1 in Beijing, China. Front. Microbiol. 5, (2014).

31. Turrientes, M. C. et al. Polymorphic Mutation Frequencies of Clinical and Environmental *Stenotrophomonas maltophilia* Populations. Appl. Environ. Microbiol. 76, 1746–1758 (2010).

32. Banar, M. et al. Global prevalence and antibiotic resistance in clinical isolates of Stenotrophomonas maltophilia: a systematic review and meta-analysis. Front. Med. 10, 1163439 (2023).

33. McCutcheon, J. G. & Dennis, J. J. The Potential of Phage Therapy against the Emerging Opportunistic Pathogen Stenotrophomonas maltophilia. Viruses 13, 1057 (2021).

34. Han, K. et al. Potential application of a newly isolated phage BUCT609 infecting Stenotrophomonas maltophilia. Front. Microbiol. 13, 1001237 (2022).

35. Han, P. et al. Biochemical and genomic characterization of a novel bacteriophage BUCT555 lysing Stenotrophomonas maltophilia. Virus Res. 301, 198465 (2021).

36. Han, P. et al. Characterization of the Bacteriophage BUCT603 and Therapeutic Potential Evaluation Against Drug-Resistant Stenotrophomonas maltophilia in a Mouse Model. Front. Microbiol. 13, 906961 (2022).

37. Li, F. et al. Isolation and characterization of the novel bacteriophage vB_SmaS_BUCT626 against Stenotrophomonas maltophilia. Virus Genes 58, 458–466 (2022).

38. Li, Y. et al. Efficacy in Galleria mellonella Larvae and Application Potential Assessment of a New Bacteriophage BUCT700 Extensively Lyse Stenotrophomonas maltophilia. Microbiol. Spectr. 11, e04030–22 (2023).

39. Zhang, W. et al. Biological characteristics and genomic analysis of a Stenotrophomonas maltophilia phage vB_SmaS_BUCT548. Virus Genes 57, 205–216 (2021).

40. McCutcheon, J. G., Lin, A. & Dennis, J. J. Characterization of Stenotrophomonas maltophilia phage AXL1 as a member of the genus Pamexvirus encoding resistance to trimethoprim–sulfamethoxazole. Sci. Rep. 12, 10299 (2022).

41. McCutcheon, J. G., Lin, A. & Dennis, J. J. Isolation and Characterization of the Novel Bacteriophage AXL3 against Stenotrophomonas maltophilia. Int. J. Mol. Sci. 21, 6338 (2020).

42. Camacho, C. et al. BLAST+: architecture and applications. BMC Bioinformatics 10, 421 (2009).

43. Nishimura, Y. et al. ViPTree: the viral proteomic tree server. Bioinformatics 33, 2379– 2380 (2017).

44. Imamovic, L. & Sommer, M. O. A. Use of Collateral Sensitivity Networks to Design Drug Cycling Protocols That Avoid Resistance Development. Sci. Transl. Med. 5, (2013).

45. Dedrick, R. M. et al. Potent antibody-mediated neutralization limits bacteriophage treatment of a pulmonary Mycobacterium abscessus infection. Nat. Med. 27, 1357–1361 (2021).

46. Bonilla, N. et al. Phage on tap–a quick and efficient protocol for the preparation of bacteriophage laboratory stocks. PeerJ 4, e2261 (2016).

47. Bushnell, B., Rood, J. & Singer, E. BBMerge – Accurate paired shotgun read merging via overlap. PLOS ONE 12, e0185056 (2017).

48. Langmead, B. & Salzberg, S. L. Fast gapped-read alignment with Bowtie 2. Nat. Methods 9, 357–359 (2012).

49. Wick, R. R., Judd, L. M., Gorrie, C. L. & Holt, K. E. Unicycler: Resolving bacterial genome assemblies from short and long sequencing reads. PLOS Comput. Biol. 13, e1005595 (2017).

50. Gurevich, A., Saveliev, V., Vyahhi, N. & Tesler, G. QUAST: quality assessment tool for genome assemblies. Bioinformatics 29, 1072–1075 (2013).

51. Brettin, T. et al. RASTtk: A modular and extensible implementation of the RAST algorithm for building custom annotation pipelines and annotating batches of genomes. Sci. Rep. 5, 8365 (2015).

52. Moraru, C., Varsani, A. & Kropinski, A. M. VIRIDIC—A Novel Tool to Calculate the Intergenomic Similarities of Prokaryote-Infecting Viruses. Viruses 12, 1268 (2020).

53. Nayfach, S. et al. CheckV assesses the quality and completeness of metagenome-assembled viral genomes. Nat. Biotechnol. 39, 578–585 (2021).

54. Alcock, B. P. et al. CARD 2023: expanded curation, support for machine learning, and resistome prediction at the Comprehensive Antibiotic Resistance Database. Nucleic Acids Res. 51, D690–D699 (2023).

55. Xie, Y. et al. TADB 2.0: an updated database of bacterial type II toxin–antitoxin loci. Nucleic Acids Res. 46, D749–D753 (2018).

56. Danecek, P. et al. Twelve years of SAMtools and BCFtools. GigaScience 10, giab008 (2021).

57. R Core Team. R: A Language and Environment for Statistical Computing. R Foundation for Statistical Computing (2022).

58. Eren, A. M. et al. Anvi’o: an advanced analysis and visualization platform for ‘omics data. PeerJ 3, e1319 (2015).

59. Kearse, M. et al. Geneious Basic: An integrated and extendable desktop software platform for the organization and analysis of sequence data. Bioinformatics 28, 1647– 1649 (2012).

60. Kutter, E. Phage Host Range and Efficiency of Plating. in Bacteriophages (eds. Clokie, M. R. J. & Kropinski, A. M.) vol. 501 141–149 (Humana Press, Totowa, NJ, 2009).

61. Gonen, S. Progress Towards CryoEM: Negative-Stain Procedures for Biological Samples. in cryoEM (eds. Gonen, T. & Nannenga, B. L.) vol. 2215 115–123 (Springer US, New York, NY, 2021).

62. Schneider, C. A., Rasband, W. S. & Eliceiri, K. W. NIH Image to ImageJ: 25 years of image analysis. Nat. Methods 9, 671–675 (2012).

63. Wickham, H., Francois, R., Henry, L., Muller, K. & Vaughan, D. dplyr: A Grammar of Data Manipulation. (2023).

64. Wickham, H. ggplot2: Elegant Graphics for Data Analysis. Springer-Verlag New York (2016).

65. Sprouffske, K. & Wagner, A. Growthcurver: an R package for obtaining interpretable metrics from microbial growth curves. BMC Bioinformatics 17, 172 (2016).

66. Blazanin, M. Gcplyr: An R Package for Microbial Growth Curve Data Analysis. http://biorxiv.org/lookup/doi/10.1101/2023.04.30.538883<x> (</X>2023) doi:10.1101/2023.04.30.538883.

